# A deep learning framework for structural variant discovery and genotyping

**DOI:** 10.1101/2022.04.30.490167

**Authors:** Victoria Popic, Chris Rohlicek, Fabio Cunial, Kiran Garimella, Dmitry Meleshko, Iman Hajirasouliha

## Abstract

Structural variants (SV) are a major driver of genetic diversity and disease in the human genome and their discovery is imperative to advances in precision medicine and our understanding of human genetics. Existing SV callers rely on hand-engineered features and heuristics to model SVs, which cannot easily scale to the vast diversity of SV types nor fully harness all the information available in sequencing datasets. Since deep neural networks can learn complex abstractions directly from the data, they offer a promising approach for general SV discovery. Here we propose an extensible deep learning framework, *Cue*, to call and genotype SVs. At a high level, Cue converts sequence alignments to multi-channel images that capture multiple SV-informative signals and uses a stacked hourglass convolutional neural network to predict the type, genotype, and genomic locus of the SVs captured in each image. We show that Cue outperforms the state of the art in the detection of five classes of SVs (including two types of complex SVs and subclonal SVs) on synthetic and real short-read data and that it can be easily extended to other sequencing platforms, such as long and linked read sequencing technologies, while achieving competitive performance. By design, Cue can also be automatically extended to support new SV classes: this versatility is crucial as novel SV types are discovered in ongoing population-scale sequencing initiatives.

## Introduction

Structural variants (SVs) are the exceptionally diverse set of all genome alterations larger than 50 base pairs. SVs encompass mutations, such as deletions, insertions, inversions, duplications, translocations, and any complex combination thereof, that can reach megabases in size. As a result, SVs account for more base-pair differences across individuals than all other variant types combined [1] and are a key driver of the genetic diversity and disease of the human genome. To date, SVs have been linked to a wide spectrum of disorders, such as cancer, autism, Huntington’s disease, Alzheimer’s, and schizophrenia [2, 3]. Although fundamental to our understanding of human genetics and advances in precision medicine, general SV discovery still remains a largely unsolved problem. This is due both to the limitations of current sequencing technologies, and, more importantly, to the challenges of effectively leveraging all the information available in the data to model and predict SVs in software, while at the same time generalizing to the wide range of SV types and sizes.

Numerous tools have been developed to date to call SVs [4, 5, 6, 7, 8, 9, 10, 11]. These methods typically extract hand-crafted features from the alignment of sequencing data to the reference genome to model both the properties of the sequencing platforms (e.g. molecule lengths, types of sequencing error, coverage statistics) and the types of SV events (e.g. mapping patterns associated with each type of SV). In whole-genome short-read sequencing, read alignment signals that are commonly used to model SVs include: *read depth* (the number of reads that map to a genome region), *discordant read pairs* (pairs of reads from the same fragment whose mapping deviates in distance or orientation from how a contiguous fragment should map), and *split reads* (reads that have several partial alignments to the reference) [12]. In long-read sequencing, *within-read discordance* and *split-read* information are typically used to detect SVs [11, 9, 10]. These signals are usually combined into a sophisticated statistical model or a heuristic rule-based pipeline to predict different SV classes and SV breakpoints. As a result, existing tools heavily rely on developer expertise and are usually tightly coupled to the properties of the sequencing data and the artifacts of preceding analysis steps (e.g. the alignment algorithm). However, given the sheer vastness of the SV landscape and the complexity of SV-informative signals, expert-driven SV detection is inherently intractable, especially for balanced SVs such as inversions and complex nested SV types, rendering us blind to major classes of genetic drivers of disease.

Deep learning offers the ability to learn complex abstractions directly from large labeled datasets without expert guidance and is hence a promising avenue for general SV discovery. Recently, DeepVariant [13] pioneered the use of deep learning for SNP and small indel calling in whole-genome sequencing (WGS) datasets. At its core, DeepVariant uses a convolutional neural network (CNN) to classify read pileup images constructed around candidate variant sites into three possible diploid genotypes and was shown to outperform all state-of-the-art variant callers and generalize well across datasets. This strategy has also been recently applied to the analysis of SVs, primarily using CNNs for SV genotyping [14, 15] and deletion detection [16]. However, while a great fit for capturing small local events (whose signature can fully fit into a single image of some standardized size), read pileup images are not well suited to capture the complex types and larger sizes of SVs. In particular, since larger SVs would not fit into one image, the following challenges arise: the joint cross-breakpoint context and signals would be lost; images that don’t overlap SV breakpoints but are internal to an SV would be challenging to classify without loss of generality; and significant complexity would be added to reconstruct one SV call from multiple separate images processed by the model. Moreover, since SVs are often nested or tightly clustered, and hence multiple SVs can appear in the same image, SV detection cannot be robustly formulated as an image classification task. This motivates the need to develop a new methodology to apply deep learning to the problem of SV calling in its full generality.

In this work, we propose a novel generalizable framework, *Cue*, for SV calling and genotyping, which can effectively leverage deep learning to automatically discover the underlying salient features of different SV types and sizes, including complex and somatic subclonal SVs. In particular, we formulate SV discovery as a *multiclass keypoint localization* task, where keypoints correspond to breakpoints of different SV type in multi-channel images. We generate input images for this task by juxtaposing two genome intervals, which can capture both SV breakpoints regardless of SV size, and simultaneously represent multiple read alignment signals as separate image channels. To perform keypoint localization, our approach employs confidence map regression using a *stacked hourglass* network [17, 18], trained to predict Gaussian response maps encoding the probability of each pixel being an SV breakpoint, allowing for multiple SVs of any type to be present in the same image, which enables our model to easily handle nested or clustered SVs. To classify SV breakpoints by type and genotype, our model outputs a confidence map for each supported SV type-genotype combination. This formulation allows Cue to be easily generalizable by design to different sequencing technologies and new SV types. In particular, to support different sequencing platforms, alignment signals encoded in each input image channel can be customized to a specific sequencing technology without requiring a change to the SV detection framework. Similarly, Cue can learn to detect new SV types directly from additional training examples that capture such SVs also without any updates to the framework. Hence, shifting to data-driven SV discovery allows Cue to keep up with and leverage numerous ongoing population-scale sequencing initiatives and sequencing technology advances.

To date, we have trained Cue to detect and genotype deletions (DEL), tandem duplications (DUP), inversions (INV), inverted duplications (INVDUP), and inversions flanked by deletions (INVDEL) larger than 5kbp; the latter two are examples of complex SVs, which have been linked to several genomic disorders [9]. To investigate the feasibility of our approach for the analysis of cancer datasets, we have also trained Cue to detect lower-frequency subclonal DELs, DUPs, and INVs. Since high-quality labeled SV callsets in real genomes are still scarce, we have trained our current models entirely on SVs modeled in silico. We have evaluated Cue using synthetic and real short-read whole-genome sequencing datasets. We show that Cue outperforms state-of-the-art methods, including Manta[4], DELLY[5], LUMPY [6], and SvABA [8], on benchmarks where ground-truth or high-confidence SV calls are available (namely, in simulation and in the HG002 GIAB Tier1 DEL benchmark[19]). To further analyze Cue’s performance on real data given the absence of additional truthsets, we have used several long-read callers (including Sniffles [9] and PBSV [10]) to orthogonally validate the results of the shortread tools. In particular, we compared the performance of short-read and long-read methods on the CHM1 and CHM13 genome mix using short-read Illumina[20] and long-read PacBio [21] datasets. We show that Cue achieves the highest relative concordance with long-read methods for CHM1 and CHM13 DELs, while lower concordance is generally observed across these technologies for the INV and DUP callsets. We analyze several examples of SVs detected by Cue and missed by other short-read tools in these benchmarks. Finally, we show a proof-of-concept extension of Cue to long-read and linked-read sequencing platforms and its performance gains, especially in complex SV discovery, as compared to several state-of-the-art long-read and linked-read SV callers. We remark that the variant types, sequencing technologies, and training data we worked with to date are just an initial set: our framework can naturally be extended to support more complex variants, additional technologies, and even combinations of technologies, which we will pursue in subsequent framework releases. The implementation of the Cue platform and the pretrained models are freely available under the MIT license at https://github.com/PopicLab/cue.

## Results

### Overview of the Cue framework

At a high-level, Cue operates in three steps: (1) read alignments are converted into images that capture multiple alignment signals across two genome intervals, (2) a trained neural network is used to generate Gaussian response confidence maps for each image, which encode the location, type, and genotype of the SVs in this image, and (3) the high-confidence SV predictions are refined and mapped back from image to genome coordinates.

To encode multiple alignment properties, we construct an *n*-channel image from alignments to two genome intervals, where *n* is the number of extracted alignment signal types. The *x*-axis and *y*-axis of this image correspond to the two genome intervals, such that a pixel maps to some range of base pairs (a locus) in each interval, and the pixel value in each channel encodes the corresponding signal extracted from the reads that map to both pixel loci (see Figure 1A). Such images capture both the local and long-range genome structure information and can uniquely characterize the type, genotype, and genomic breakpoints of an SV. Note that the pixel corresponding to the start of the SV on the *x*-axis and the end of the SV on the *y*-axis simultaneously encodes both breakpoints (we refer to such pixels as *breakpoint keypoints*). Therefore, by juxtaposing intervals that are close-by and distant on the genome, we can depict breakpoints of both small and large SVs in the same image. Using a streaming sliding-window approach to scan the genome, we produce candidate genome interval pairs and the corresponding images on the fly.

**Figure 1:**
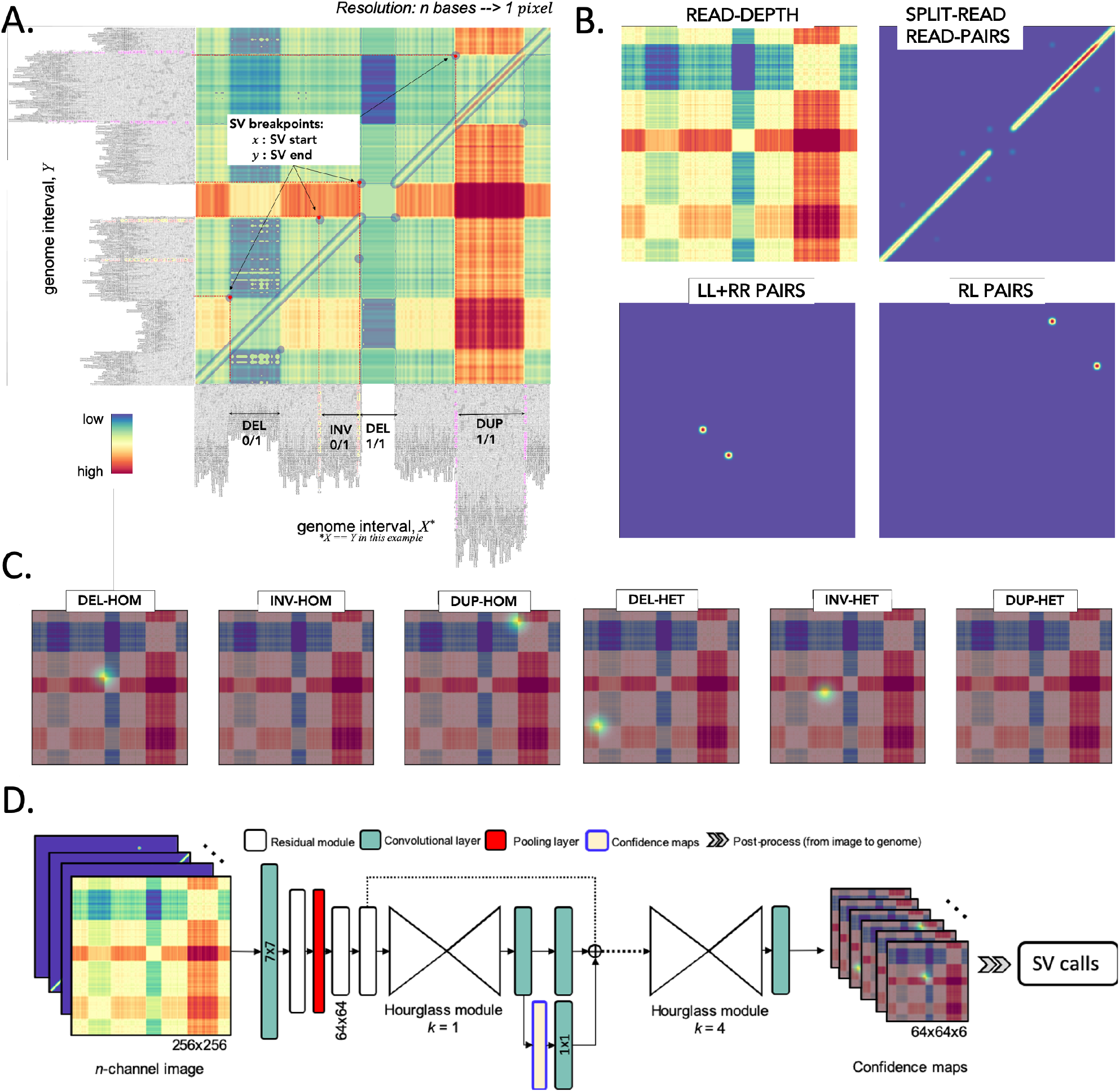
Overview of the Cue framework. **A**. Conversion of sequence alignments to images. Alignments from a genome interval (visualized in IGV[22]) are shown on the *x*-axis and *y*-axis of the resulting image (displaying the overlay of several signal channels), annotated with four different SVs in this interval. The four highlighted pixels (keypoints) in the image correspond to the breakpoints of each SV (given by their start coordinate on the *x*-axis and their end coordinate on the *y*-axis). **B**. SV image channels representing different signals. **C.** SV breakpoint confidence maps (predicted by the network for the image in **A**) for the following SV types and genotypes: homozygous (HOM) and heterozygous (HET) deletions (DEL), inversions (INV), and tandem duplications (DUP). For simplicity only the read-depth channel is shown as the background. The bright kernels in each map represent a high confidence that the breakpoints of an SV of that specific type occur at that location, e.g. each pixel in the DEL-HOM map encodes its probability to be a homozygous deletion keypoint. **D.** The architecture of the stacked hourglass network used in Cue. It takes an *n*-channel image as input and generates a confidence map for each supported SV type. The predicted confidence maps are then post-processed to produce the final SV callset.

We encode the following alignment signals from short reads: read-depth, split-read, read-pairs, and the discordant read-pair orientations – right-left (RL), left-left (LL) and right-right (RR) – where LL and RR pairs are indicative of an inversion and RL pairs signal a duplication. We use different functions to represent these signals (or their combination) in each corresponding channel as described in the Methods section. For example, readdepth signal is computed as the difference in depth between two loci. Figure 1B shows several image channels, visualized as heatmaps, generated from read alignments to a 150kbp synthetic genome interval. Briefly, in addition to read-depth: the split-read/read-pair channel shows the number of split reads or read pairs mapping to both loci; the LL+RR read-pair channel shows the number of read pairs that map to both loci in the same orientation (such as a forward-forward or reverse-reverse mapping), and the RL read-pair channel shows the number of read pairs that map to both loci in the RL orientation (where the second read in the pair maps to an earlier position on the reverse strand and the first read of the pair maps to a later position on the forward strand). We stack the resulting heatmaps into an *n*-channel image and use this image as input to our deep learning model. In order to support new sequencing platforms, we can reuse and extend the library of alignment signals and functions used to generate the input channels. Furthermore, by combining channels generated for multiple sequencing platforms into the same input image, we can also automatically train our framework on inputs derived jointly from multiple platforms. Finally, given the well-known association between SVs and repeats (see e.g. [23]), we could design additional channels to make our calls more robust in highly repetitive regions (e.g. by encoding all the known repeats in the reference). These extensions will be addressed in future work.

The resulting images can contain multiple SVs of different type and size, as well as partially visible SVs (captured when the genome intervals overlap but do not fully include an SV). Moreover, unlike physical real-world objects, which are self-contained and separable from each other and their background (having explicit boundaries), SVs form patterns, which interact in intricate ways and don’t have a well-defined boundary. To address these challenges, we formulate SV detection in these images as a *multi-class keypoint localization* task, wherein the objective is to detect the image coordinates corresponding to the two breakpoints of each SV in the image (i.e. the breakpoint keypoints), categorized by SV type and genotype. We solve this task using confidence map regression by training our network to predict a set of (Gaussian response) confidence maps for each image, such that each map corresponds to a zygosity-aware SV type supported by the model (e.g. heterozygous deletion or homozygous inversion) and encodes the location of the breakpoints of all SVs of this type in the input image. Figure 1C shows the set of six predicted confidence maps, used to detect deletions, inversions, and duplications (split by genotype). To create ground-truth confidence maps for an image (given a ground-truth BED or VCF file), we place an unnormalized 2D Gaussian kernel centered at each breakpoint keypoint in the confidence map given by the type and genotype of each SV represented in the image. Given this approach, we can easily generalize our framework to detect a new SV type by configuring it to predict two additional confidence maps (for both genotypes) and just feeding it new examples labeled with this SV type.

To generate accurate confidence maps, our neural network needs to leverage features at both local and global scale to learn the structural complexity of SV signatures and multi-SV interplay patterns. To this end, Cue uses a fourth-order stacked hourglass convolutional neural network (HN) based on [17, 18], which can consolidate information at multiple scales by repeated bottom-up (pooling) and top-down (upsampling) processing and intermediate supervision. Figure 1D depicts the high-level network architecture of the HN model used by Cue. The network takes an *n*-channel image as input and outputs a set of confidence maps, encoding the breakpoint keypoints of all SVs in the image (split by type). The mean squared error or L2 loss is commonly used to measure the distance between the predicted and the ground truth confidence maps. However, confidence maps encoding a few keypoints using Gaussian kernels mostly consist of background pixels (of value zero), which creates a severe imbalance between foreground and background classes. To address this imbalance, we use *focal L2 loss* adapted from [24], which allows us to scale down the contribution of easy background and easy foreground pixels when training Cue (see Methods).

Finally, given the confidence maps regressed by the network, we produce the set of final SV calls as follows: (1) we detect all local peaks in each confidence map, (2) refine the keypoint positions using high-resolution images, and (3) convert the refined keypoints to genome space to obtain the genome SV breakpoint coordinates. Note that the type and genotype of each SV are directly given by the respective confidence map indices. In addition, non-maximum suppression (NMS) filtering of lower-confidence conflicting calls is performed in both image (2D) and genome (1D) space. Details about each step of the Cue framework are provided in the Methods section.

### DEL, DUP, and INV discovery from short-read synthetic data

To benchmark Cue in single event detection, we simulated a human genome using SURVIVOR[25] based on the GRCh38 reference with a total of 13,504 SVs of size 5kbp – 250kbp, with the following breakdown by type: 4,500 DELs, 4,461 DUPs (tandem), and 4,543 INVs. We simulated a large number of SV events to create a more challenging benchmark, which guarantees that some SVs are placed in difficult regions of the genome (e.g. segmental duplications) and clustered near each other. To evaluate the effect of the genome sequence context on SV calling, we categorized the simulated SVs based on their position into the following four types (defined in [26]) using the RepeatMasker[27] and segmental duplication tracks from the UCSC genome browser[28]: (1) segmental duplication (SD), (2) simple repeat (SR), (3) repeat masked (RM; all other repeats excluding SD and SR), and (4) unique. The resulting breakdown of the simulated SVs by context type is as follows: SD=1,131(8.3%), SR=786 (5.8%), RM=8,722 (64.6%), and unique=2,865 (21.2%). We then simulated a 60x paired-end Illumina short-read WGS dataset using DWGSIM [29] from this genome and mapped the reads with BWA-MEM[30] to obtain the input to the methods evaluated in this benchmark.

We compare the performance of Cue in calling and genotyping the simulated events above to four popular state-of-the-art SV callers, namely: Manta [4], LUMPY [6], DELLY [5], and SvABA [8]. Figure 2A shows the precision, recall, and F1 score of each method in calling and genotyping SVs computed using the benchmarking tool Truvari[31] (see Methods for more details on evaluation metrics and the Supplementary Note for tool execution details). As shown in Figure 2A and Supplementary Figure A1, Cue consistently achieves the highest scores in the three reported metrics when calling SVs, leading by 1-15% in F1 score and 2-12% in recall. Manta and LUMPY achieve equally high precision across all SV types, with a 2-6% loss in recall depending on the SV type. When genotyping SVs, Cue achieves the highest scores in all the metrics on average across all SV types, with a gain in F1 of 5-54%. The biggest increase in F1 score is seen for genotyping DUPs, where Cue leads by 12-36%. On the other hand, Manta and LUMPY outperform Cue by 1% in F1 when genotyping INVs. The performance profile (i.e. the precision vs recall trade off) of DELLY and SvABA varies most considerably in this benchmark both in calling and genotyping SVs of different types, although slight variation can be observed for most tools. To further examine the effect of SV type, size, and genome context on the performance of each tool, Figure 2B shows each method’s false negative (FN) calls broken down by size, type and genome context. We can see that method performance can vary significantly depending on these features. For example, most tools miss events of smaller size that fall into SD and SR regions of the genome. Cue misses the fewest events in such repeat regions – namely, 171 SD events and 172 SR events. LUMPY achieves the second highest recall, missing 408 SD events and 258 SR events.

**Figure 2:**
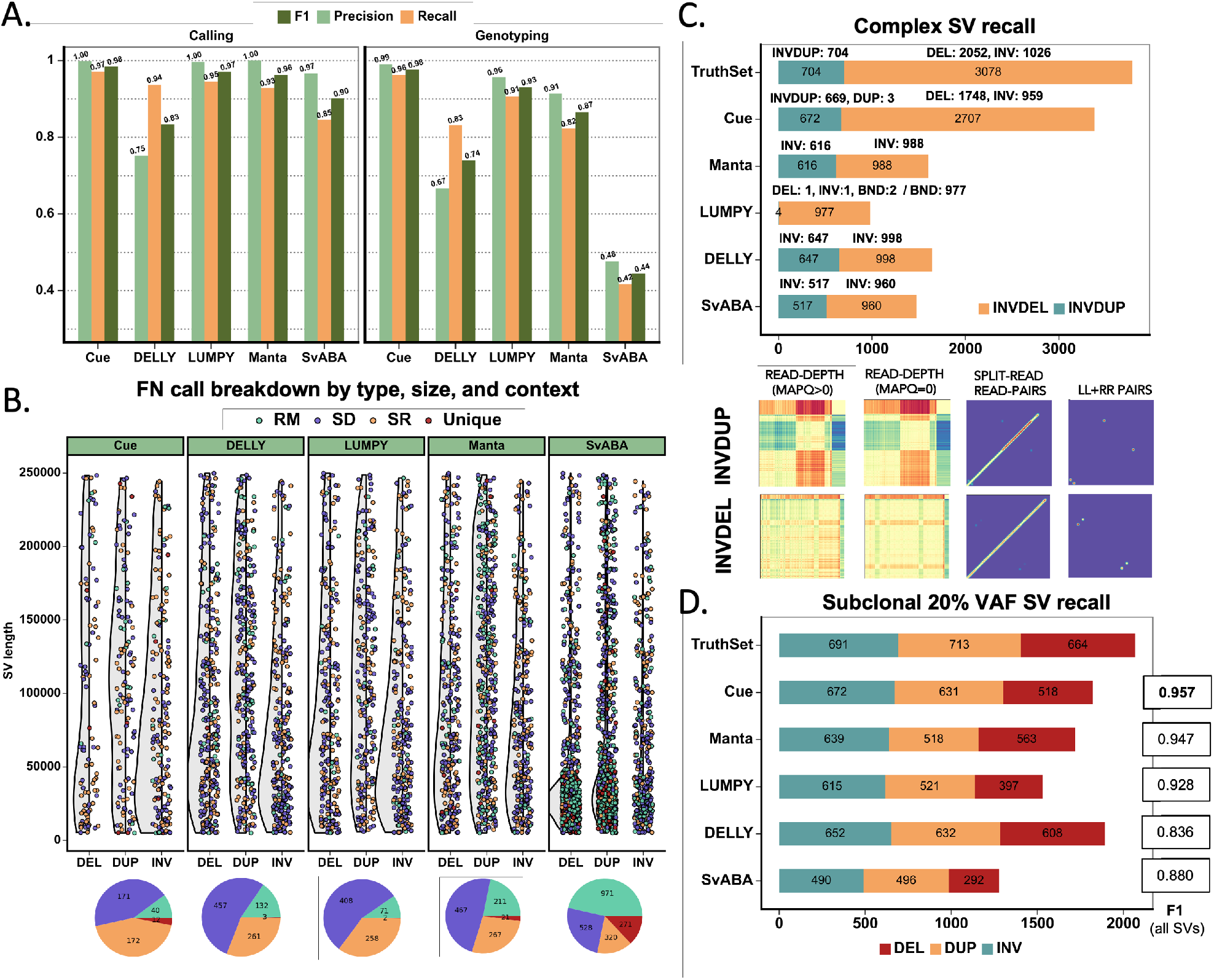
Performance evaluation on synthetic data. **A.** Recall, precision, and F1 in SV calling and genotyping on 60x paired-end short-read WGS data. **B.** FN calls broken down by size, SV type, and genome context. **C.** Recall of two complex SV types (INVDUP and INVDEL). Bottom: examples of each event type found only by Cue. **D.** Recall of subclonal somatic SVs broken down by type.

### Complex SV discovery from short-read synthetic data

Since our deep learning framework is designed to detect the breakpoints of any number of SVs in the same image, it can be naturally leveraged to detect clustered and complex SVs. To that end, we trained Cue to additionally detect the following two complex SV types: deletion-flanked inversions (INVDEL) and inverted duplications (INVDUP). We represented INVDUPs in Cue as a separate SV type, while detecting INVDELs as three separate SVs (two DELs and one INV). For this benchmark, we simulated a human genome using SURVIVOR with 7,240 SVs of size 5kbp – 250kbp in total, of which 1,048 were INVDELs and 704 were INVDUPs. We configured Truvari to count a variant as a true positive regardless of its reported type so as not to penalize tools that do not specifically detect or label complex subtypes. Figure 2C shows the recall of complex SVs of each tool broken down by type, as well as an example of how each event type is captured across several image channels constructed by Cue. Cue discovered a significantly higher number of complex events in this benchmark; in particular, it found 88% of INVDELs, which is >55% greater than the next best result (32% found by DELLY), and 95% of INVDUPs. Of the discovered INVDUPs, Cue labeled over 99.6% events correctly as INVDUPs (with only 3 events called as DUPs). Manta and SvABA reported all recovered INVDUPs as INVs; DELLY reported INVDUPs as INV events; and LUMPY detected only 4 INVDUP events reported as a DEL, INV, and two breakends (BNDs). For INVDELs, Manta, SvABA, and DELLY did not call any flanking deletions, while LUMPY called all matching events as BNDs. Supplementary Table A1 additionally shows the percentage of recalled variants with a correct genotype for each tool. Cue assigned the correct genotype to >98% of all discovered events. It significantly outperformed other tools in genotyping INVDUPs, where SvABA achieved the largest percentage of correctly genotyped discovered INVDUPs of 68%. Finally, since Cue reports INVDUPs directly, we could evaluate its precision as well and found that Cue does not report any false positive INVDUPs, achieving 100% precision for calling INVDUPs in this benchmark.

### Subclonal somatic SV discovery from short-read synthetic data

Somatic SV discovery in cancer genomes is complicated by tumor heterogeneity, wherein some SVs may be present only in certain tumor subclones corresponding to a fraction of the reads in the dataset. As a result, such SVs will have lower variant frequencies (VAFs) and can be challenging to distinguish from noise using manually-designed heuristics. Here we investigate the ability of our method to automatically learn to detect lower VAF subclonal SVs, which will produce fainter signals in our images, using synthetic data.

In order to generate WGS data with subclonal SVs, we first simulated two human genome haplotypes (A and B) using SURVIVOR based on the GRCh38 reference. We then created a FASTA file with one copy of haplotype B and four copies of haplotype A, and simulated a paired-end short-read dataset with a total coverage of 60x from this FASTA file as before using DWGSIM. This resulted in an effective 20% VAF for SVs of haplotype B. We used half of the chromosomes of this genome to train Cue, and the other half for evaluation. The resulting evaluation SV callset had a total of 6,266 SVs (heterozygous and homozygous events across both haplotypes) and 2,068 subclonal SVs from haplotype B. Figure 2D shows subclonal SV recall results broken down by SV type, as well as the F1 score achieved by each method of the full evaluation dataset. Cue recovered 97% of all subclonal SVs (96% of deletions, 96% of duplications, and 98% of inversions) while maintaining the highest F1 score, and outperformed other methods in subclonal INV discovery. While DELLY recovered the most DEL and DUP events, it also achieved the lowest F1 score (12% lower than Cue) in this benchmark.

### HG002 GIAB DEL benchmark

To evaluate Cue on real data, we used the HG002/NA24385 sample and its hg19 GIAB NIST Tier1 v.06 benchmarking callset [19] available from the Genome in a Bottle (GIAB) Consortium. This callset contains curated deletion and insertion calls obtained by consensus calling with multiple sequencing technologies; we used only calls from the high-confidence regions defined in this release. We obtained the 60x Illumina HiSeq short-read BAM from GIAB (and re-mapped the reads using BWA-MEM) and evaluated Cue and other tools using this dataset. Figure 3A shows the performance in SV calling obtained by the five methods computed using Truvari for deletions greater than 5kbp. In this benchmark, Cue outperformed other methods by 4-23% in F1 score. Manta achieves the highest precision (2% greater than Cue); however, its recall is 11% lower than Cue. On the other hand, LUMPY achieves the highest recall (1% greater than Cue) with a precision drop of 6% compared to Cue. Figure 3B shows the false negative (FN), false positive (FP), and true positive (TP) DEL calls broken down by size and genome context with several highlighted events, for which IGV plots and Cue-generated image channels are shown in Figure 3B and Supplementary Figure A2. In particular: (1) is a TP LINE-1 (L1HS) deletion event detected only by Cue, (2) is a FP event called by DELLY, LUMPY, and Manta, and not called by Cue, and (3) is a FP event called by DELLY, LUMPY, Manta, and SvABA, and not called by Cue. As we can see in the Cue-generated channels, the LINE-1 deletion signature of event (1) is well-captured in the high-MAPQ read-depth channel (which shows the drop in coverage consistent with a deletion or a repeat) and the split-read/read-pair channel (which shows the novel adjacency formed by discordant reads-pair mappings). The signatures in these two channels jointly, along with the absence of signal in the remaining channels, can uniquely characterize a deletion of a repeat element. On the other hand, while the split-read/read-pair channel alone at the site of FP event (2) can be consistent with a deletion, the presence of the RL signal and the absence of read-depth signal are not jointly consistent with a deletion. We found that the mapping signature observed for the FP event (2), along with several variations, is commonly reported as a DEL event by short-read SV callers, or sometimes as two separate DUP and DEL events. While, theoretically, a precisely overlapping DEL and DUP event on each haplotype can produce a similar mapping pattern, additional properties of this mapping signature reveal that they are often the result of either a dispersed DUP or a divergent reference repeat (defined as the presence of two inexact copies of a locus in the reference sequence) as illustrated in Supplementary Figure A3. In particular, we performed an in-depth analysis of the loci at the breakpoints reported for event (2) using PacBio CCS reads, which revealed it to be a divergent repeat in the reference. More specifically, from the joint de Bruijn graph of the reference region and of the CCS reads that were mapped to the region, we found that there are two copies of a short sequence in the donor genome (approximately 500bp around the two breakpoints), where one copy (on the right) matches the reference, and the other copy involves several mismatches with the reference (on the left). A pairwise global alignment of the two sequences in the reference (computed using the Needleman-Wunsch implementation in EMBOSS Needle [32]) revealed a 92% identity, while the corresponding sequences extracted from CCS reads from the donor had a 99% identity. The short-read mapping patterns further support this hypothesis, with a consistent gain and loss of coverage observed at the two breakpoints and a discrepancy in the span of the RL and large-insert read pairs (green and red pairs shown in the IGV plot) consistent with the schematic in Figure A3. Finally, similar to event (2), the FP event (3) displays no drop in read depth, and a complex pattern of discordant read pairs that cannot be explained by a single DEL. Validation with PacBio CCS reads shows that this signature is induced by two dispersed DUPs and one inverted dispersed DUP (see Figure A2 for details). Directly learning from labeled positive and negative examples, Cue is able to leverage the information across all the channels jointly to make accurate predictions for these events, and in this instance it can correctly learn that read pairs with a discordantly large insert size are not enough evidence by themselves for calling a DEL.

**Figure 3:**
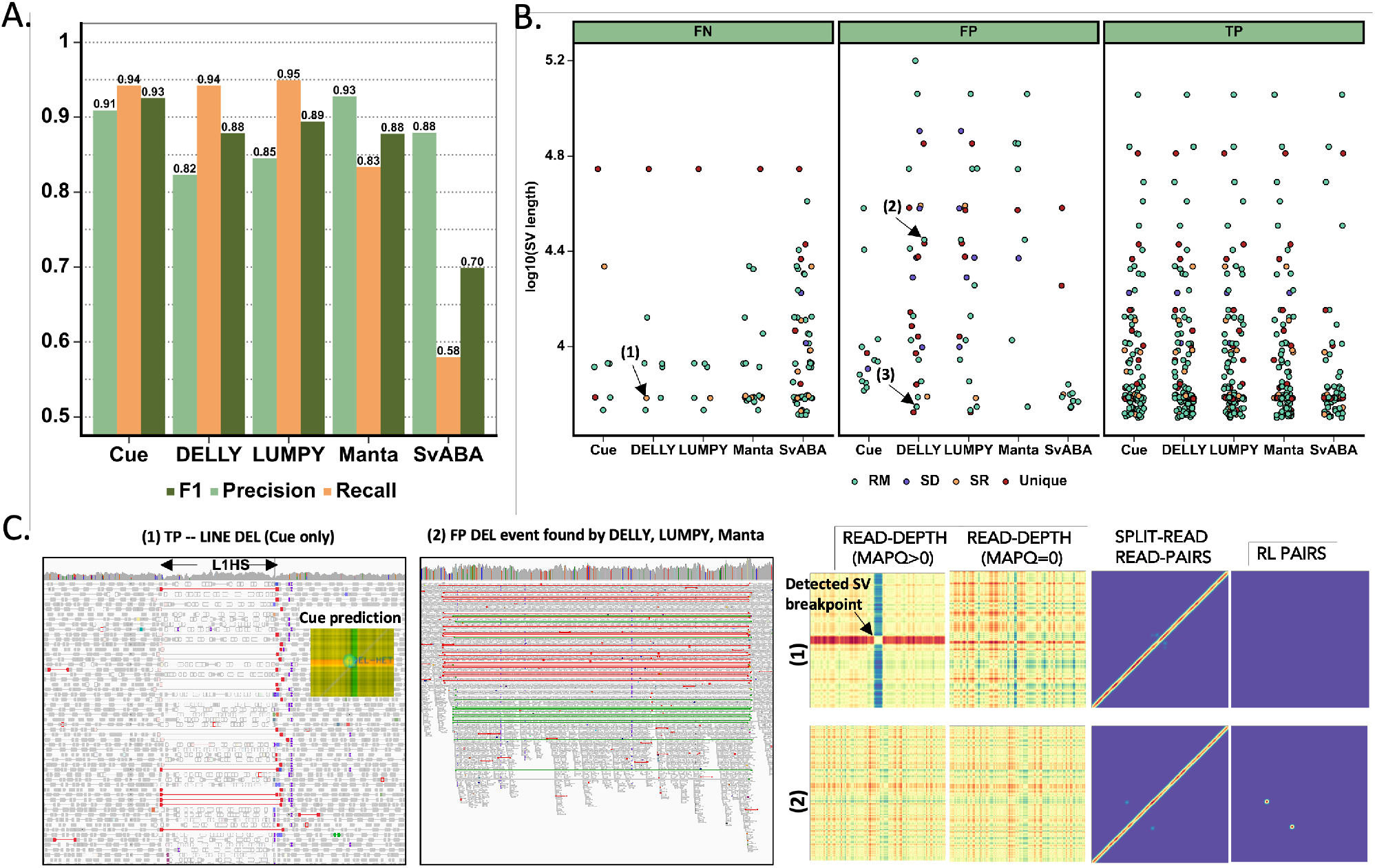
Performance evaluation on HG002 GIAB deletion benchmark. **A.** Recall, precision, and F1 in DEL calling. **B.** FN, FP, and TP calls broken down by size and genome context. (1)-(3) are FN and FP SV calls analyzed in panel C and Supplementary Figure A3. **C.** IGV plots and Cue image channels for a TP and FP call: (1) TP LINE-1 deletion event detected only by Cue and (2) FP deletion call made by DELLY, LUMPY, and Manta (involving a diverging repeat in the reference illustrated in Supplementary Figure A2). IGV shows RL read-pair alignments in green and read pairs with a discordantly large insert size in red.

### CHM1 and CHM13 diploid mix benchmark

We further evaluated Cue using the haploid hydatidiform mole CHM1 and CHM3 cell line samples. We obtained the Illumina WGS reads for each sample (at 40x coverage each), merged the reads to create a diploid 80x coverage mix in silico, and mapped the resulting mix against the GRCh38 reference using BWA-MEM. Since a high-confidence truthset is not available for these two genomes, we used three callsets derived from PacBio CLR long reads to orthogonally validate the results of the short-read SV callers. In particular, we used the published Huddleston et al. callset[21] and generated calls from two other long-read SV callers, Sniffles[9] and PBSV[10], on the PacBio CHM1 and CHM13 long reads published by [21]. Figure 4A is an upset plot depicting the agreement across all evaluated callsets for DEL events larger than 5kbp. Tool callset overlaps were computed using SURVIVOR (see the Methods section for details). All long and short read callers discovered the same 82 DEL events, with an additional 34 events found by everyone except SvABA. Of the short-read callers, DELLY and LUMPY have produced the largest set of unique calls (i.e. calls that were not reported by any other tool), 264 and 105, respectively. Cue has produced 49 unique calls and Manta has produced the fewest unique calls (only 9). Since these events are likely to be FPs, we can estimate that DELLY and Manta achieved the lowest and highest precision, respectively, on this dataset, which is consistent with the HG002 benchmark. In order to estimate recall, we looked at how many short-read calls were validated by at least one long-read caller. As seen in Figure 4B, Manta and SvABA had the lowest number of reported calls also discovered using long reads, while LUMPY and DELLY had the highest, which is also consistent with their performance profile on the HG002 benchmark. We can see here that the significant majority of Cue calls (186 out of 244) were found by at least one long-read method – the largest fraction of calls of all short-read methods. Of these calls, 7 were found only by Cue, Sniffles, and Huddleston et al. Next we manually examined several groups of events that were found by multiple short-read callers only (i.e. they lacked long-read validation). For example, 68 DELs were reported by DELLY, LUMPY, and SvABA, while 42 events were reported by all short-read callers except Cue. Even though tool consensus is high for these sets of events, we found that a large fraction of them appear to be likely false positive calls caused by a dispersed DUP event or a divergent repeat in the reference, similar to false positive DEL call signatures reported for HG002. For example, 12 of the 42 events had a DUP call reported by at least one of the four methods at the same locus. Figure 4C shows the IGV plot of such a DEL call found by DELLY, LUMPY, and SvABA, while Supplementary Figure A4A shows such a DEL call found by all four short-read methods except Cue. Figure 4C also shows an example of a true LINE-1 DEL event found only by Cue and two long-read methods and Figure A4B shows a similar LINE-1 DEL event found only by Cue and all the long-read methods. Consistently with HG002 results, Cue is able to detect some LINE-1 DEL events missed by all other short-read callers and validated by long-read callers in this benchmark as well.

**Figure 4:**
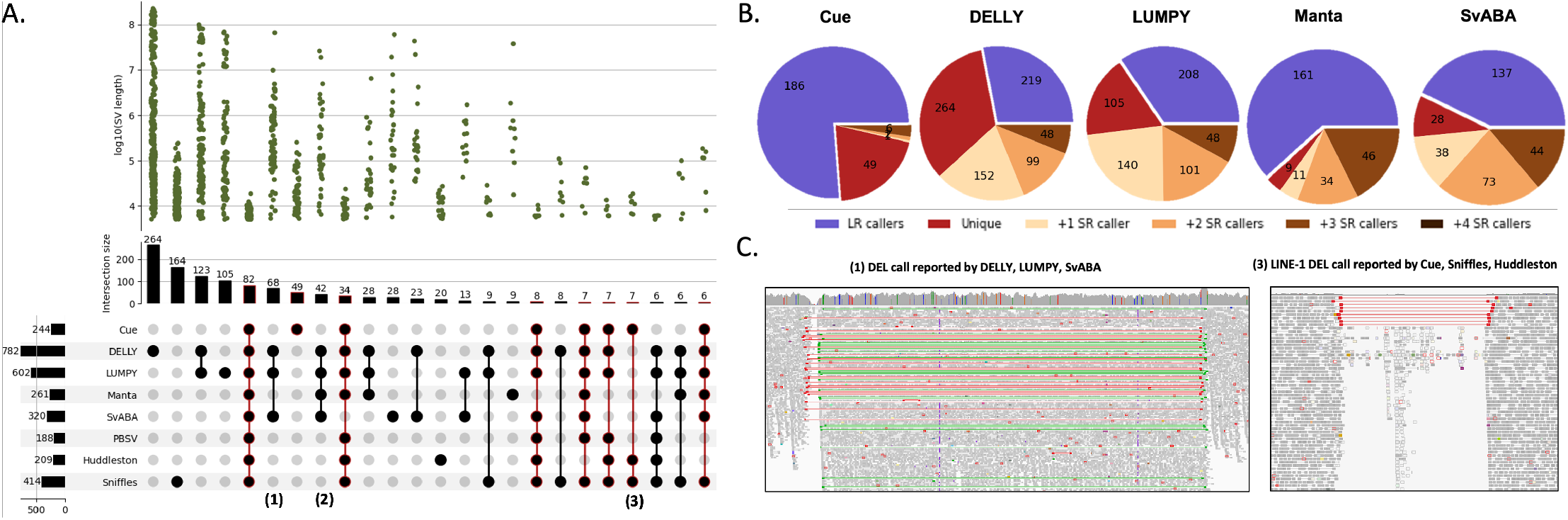
Performance evaluation of DEL calling on the CHM1+CHM13 cell-line mix. **A.** Upset plot depicting DEL callset overlaps of five short-read and three long-read callers (only sets larger than 5 events are displayed for conciseness). **B.** Result breakdown by consensus with long-read and other short-read callers. **C.** IGV plots of (1) a DEL event reported by DELLY, LUMPY, and SvABA and (2) a LINE-1 DEL detected by Cue and long-read callers only.

We performed a similar analysis for DUP and INV events reported in Supplementary Figure A5. As opposed to the DEL benchmark, the consensus across callers is considerably smaller for these event types. In particular, only one DUP and zero INV events were consistently reported by all of the short and long-read callers. As can be seen in Supplementary Figure A5A-B, all short-read methods agreed on 10 DUP events (with no long-read support) and 7 DUP events were consistent with Sniffles alone. A considerable number of events were reported by multiple short-read callers except Cue. In particular, 32 DUPs were found by Manta, LUMPY, DELLY, and SvABA. Since consensus for these calls was high, we manually examined each call individually. We found that 18 out of the 32 events were found within centromeric satellite repeat regions of multiple chromosomes. Since these regions are dominated by repeats, they are notoriously difficult to analyze using short reads. As can be seen in Supplementary Figure A6, RL signatures are heavily present in these regions and can potentially lead to FP DUP calls by methods that rely only on such mappings to identify DUP events. For example, we can see that the methods produce numerous (overlapping and inconsistent) DUP calls for the displayed genome region on chr12. Of the remaining 14 calls, we found that 8 were also reported as DEL events by some of the tools and were consistent with either the dispersed DUP or the divergent reference repeat signature discussed above. As seen in Supplementary Figure A5C-D, the overlap of INV callsets is substantially smaller across sequencing technologies. In particular, all short-read methods except LUMPY agreed on only 13 INV events (with no long-read support) and another 13 INV events were found only with long-read methods. Overall these results point to the difficulty of calling DUPs and INVs using existing methods, and using orthogonal technologies for validation. More importantly, they reveal the need for a benchmarking dataset for such events (and especially complex SVs) that can be used to evaluate existing and future tools.

### Extending Cue to long and linked read sequencing platforms

To demonstrate that our approach generalizes to different sequencing platforms, we have also performed a preliminary evaluation of Cue on long-read and linked-read data. To leverage longer read lengths, we have added a new alignment signal channel, the *split-molecule* channel, which captures the number of long molecules spanning two given loci. In linked reads, the split-molecule signal can be readily extracted by counting the number of barcodes shared by reads aligned to each locus. In order to extract the split-molecule signal (and the read-depth signal) from long reads, we first generate a short-read coverage profile for each long read (wherein short reads are extracted from the long read, barcoded uniquely with their long read name, and remapped to the long read alignment window). The short-read coverage profile allows us to easily capture within-read SV events (e.g. duplications and inversions) without sacrificing long-read mappability. Supplementary Figure A7A shows the image channels generated from PacBio CLR long reads and 10x Genomics linked reads, respectively, for the genome interval in Figure 1A. For long reads, we have also included the clipped-read signal computed as the difference in clipped-read coverage at two loci, along with the split-read and read-depth channels. For linked reads, we have directly leveraged all the read-pair channels developed for short-read paired-end sequencing. We have simulated long reads at 30x coverage using PBSIM2 [33] (aligned with minimap2 [34]) and linked reads at 60x coverage using LRSIM [35] (aligned with Longranger [36]) from the synthetic training and evaluation genomes used in our short-read benchmarks.

We compared the performance of Cue (i.e. of the models trained only on linked reads and only on long reads, respectively) to two state-of-the-art long-read SV callers, PBSV [10] and Sniffles [9], and to two linked-read SV callers, Longranger [36] and LinkedSV [37]. We performed the evaluation on the two synthetic benchmarks described above for single-event (DEL, INV, and DUP) and complex-event (INVDUP and INVDEL) discovery. As shown in Figure A7B, Cue achieves the highest scores in DEL, INV, and DUP calling with these sequencing technologies. We see the biggest performance gap in comparison to linked-read methods, where Cue leads by 15% in F1 score. Note that state-of-the-art linked-read callers are also outperformed by existing *short-read* callers: this suggests that developing expert-driven software that fully leverages linked-read signals is particularly difficult, making this data type an ideal application domain for deep learning methods. Cue achieves significantly higher genotyping accuracy with both long and linked reads as well.

Figure A7C shows the recall of complex SVs of each tool, broken down by type. As with short reads, Cue discovered a significantly higher number of complex events in this benchmark than existing approaches. In particular, it found 98% and 97% of INVDUPs using long reads and linked reads, respectively, while only 56% and 64% of these events were found by the leading tools, PBSV and Longranger, respectively. Cue labeled 100% of the detected events correctly as INVDUPs using long reads and 98% using linked reads. Existing methods reported most of the recovered INVDUPs as INVs. Of the existing tools, only Sniffles was designed to detect INVDUPs specifically; however, of the recalled 382 (54%) SVs, it reported only 7 as actual INVDUPs, and the majority as INVs. Furthermore, Cue found 83% (with long reads) and 84% (with linked reads) of INVDELs, respectively, which is significantly greater than the next best result of 34% found by Sniffles. For INVDELs, existing methods missed most of the flanking deletions and discovered only a fraction of the inversions.

In conclusion, by just applying minimal modifications to the image channels, our approach can be extended to other, completely different sequencing platforms, while matching or surpassing state-of-the-art methods that were manually tailored to those platforms. This combination of power and simplicity of deployment allows Cue to keep up with the rapid advances in sequencing technology and to deliver optimal performance on each platform.

## Discussion

In this work we motivate the use of deep learning for structural variant discovery, which allows us to shift method development away from ad hoc hand-engineered models and heuristics-based pipelines to scalable and sustainable models that can learn complex patterns of variation automatically from the data. We lay out how SV detection can be formulated as a deep learning computer vision task and propose a novel framework, Cue, to call and genotype SVs of diverse size and type. We demonstrate state-of-the-art results in calling several SV classes, including complex and subclonal SVs, from synthetic and real short-read datasets. To date, our model was trained to detect both simple germline and subclonal deletions, inversions, and tandem duplications, as well as complex deletion-flanked inversions and inverted duplications, larger than 5kbp. By design, Cue can be easily generalized to other sequencing technologies (such as PacBio, Oxford Nanopore, stLFR, and Hi-C), and combinations thereof, by adapting or expanding the set of alignment signals used to generate the images, without requiring any other updates to the framework. For a proof of concept, we have adapted our framework to PacBio CLR long reads and 10x Genomics linked reads and show that Cue can outperform existing methods on these platforms, especially in complex SV discovery. Even more crucially, since Cue learns directly from the data, it can be easily trained to detect overlapping SVs from different haplotypes, as well as new complex SV types as they are discovered by the genetics community, provided that we have enough labeled examples at hand. However, since it relies on read alignment signals, Cue can detect just the breakpoints of novel insertions, and needs to be integrated with a local assembler to reconstruct the inserted sequence itself. We will expand the framework in future work with support for additional sequencing platforms, smaller SV sizes (which can be achieved by changing the basepair-to-pixel resolution of our images), and more SV types (such as translocations, insertions, and other complex SVs).

A major requirement and challenge for data-driven SV discovery is the availability of large and well-balanced training datasets. While numerous callsets are available in publicly-available repositories, such as GIAB and HGSVC [38], high-confidence calls that can be used reliably for training are currently very scarce and are mostly composed only of simple deletion and insertion events (e.g. the GIAB HG002 truthset used here for benchmarking). Note that using high-confidence calls of just a subset of event types is challenging for training without manual validation, since such calls may fall in the proximity of undetected SVs of a different type and would effectively label such events as negative examples. To compensate for the current lack of available real-genome training data, we have used in silico SV modeling for training. This approach can produce arbitrarily large well-balanced datasets; however, extensive modeling and parameter sweeps are required to capture the full repertoire of sequencing technology characteristics, SV types, and genome contexts that we expect to see in real data. While training on synthetic data produced encouraging results, we expect the model to struggle with event types it has never seen during training; therefore, including real data into the training dataset is critical both for performance and generalizability. Importantly, we can keep up with the rapid growth of available sequencing datasets and our understanding of structural variants, by retraining our model on new data as our SV truthsets grow over time, while leaving our core framework untouched.

## Methods

### Multi-channel image generation

Given a set of read alignments *A* from an input BAM file, a set of candidate genome interval pairs *G* (with intervals of size *S*), and a set of *n* predefined alignment signal scalar valued functions *F* = {*f*^1^, *f*^2^,…, *f^n^*} (where 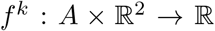) mapping alignments extracted from two regions of the genome to a single value, we create an *n*-channel *w* × *h* image for each interval pair (*g_x_*,*g_y_*) ∈ *G*, where the *k*th image channel is obtained as follows. First we use the function *f*^k^ ∈ *F* to compute a square matrix **M**^*k*^, such that 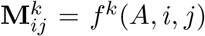. The dimension of **M**^*k*^ is 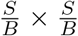, where *B* represents the matrix *resolution* (or the number of genome base pairs that correspond to one entry in **M**). The *i^th^* row in **M**^*k*^ corresponds to the genome region (or bin) *g_y_*[*iB*…(*i* + 1)*B*), the *j*^th^ column corresponds to the genome region *g_x_*[*jB*…(*j* + 1)*B*), and the value 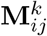. is given by applying *f^k^* to alignments in regions *i* and *j* (denoted as the alignment subsets *A^i^* and *A^j^*, respectively). We assign an alignment to a region if its midpoint position falls into the region. Post-construction, each matrix is down-sampled (using block summation) or up-sampled (using nearest neighbours) to size *w* × *h*, depending on the configured values of *S*, *B*, *w*, and *h*, and its values are normalized to fall in the range [0, 1]. In the resulting image, genome positions from the interval *g_x_* are on the *x*-axis and those from *g_y_* are on the *y*-axis, respectively. Presented results were obtained with images of size 256 × 256 pixels, *B* of 750bp for SV discovery and *B* of 200bp for SV refinement, and *S* of 150kbp.

We define the following alignment signal functions (note: we denote the sets of a specific property taken from all elements in a given alignment set using subscript notation. For example, the set of all read names taken from the alignments in a subset *A^i^* is given by 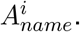):

- The read-depth function

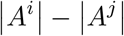

computes the difference in coverage in regions *i* and *j* normalized such that negative values fall in the range [0, 0.5), and positive values fall in the range (0.5, 1] (to distinguish between deletion and duplication events); we compute two channels using this function with (1) only MAPQ>0 alignments and (2) all alignments (including MAPQ=0).
- The split-read and read-pair function

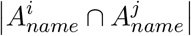

computes the number of reads or read pairs mapping to both bin *i* and *j*, where read pairs and split-read alignments are given the same name.
- The read-pair LL and RR orientation function

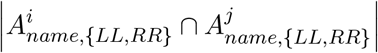

computes the number of read pairs that map to both bin *i* and *j* in the same orientation, such as a forward-forward or reverse-reverse mapping; these mappings are common with inversions.
- The read-pair RL orientation function

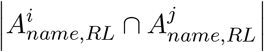

computes the number of read pairs that map to both bin *i* and *j* in the RL orientation, where the second read in the pair maps to an earlier position on the reverse strand and the first read of the pair maps to a later position on the forward strand; these mappings are common with duplications.
- The read-pair orientation function

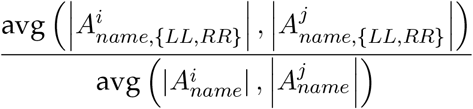

computes the ratio in average coverage by LL and RR read pairs versus all pairs, which is indicative of the inversion genotype.

In order to boost the signal from discordant mappings, the normalized output of the read-pair orientation functions is also passed through a dilating maximum filter and a Gaussian filter.

To speed up computation, we construct an index of the input BAM file that allows us to quickly query several properties of the alignments mapped to individual genome bins. More specifically, we partition each chromosome into *B*-sized bins, and store the following sets of values extracted from the reads assigned to each bin: (1) number of MAPQ>0 reads, (2) number of all reads, (3) read names, (4) read names of the LL and RR read-pairs, (5) reads names of the RL read-pairs. Each function can then be easily computed via index lookups (e.g. the split-read and read-pair function can be computed as the intersection of the read names stored in the corresponding two bins of the index).

### Interval pair selection

We use the following sliding-window strategy to generate *G*, the set of candidate interval pairs (for which images are constructed) that capture both small and large SV events on each chromosome. Let *L* be the length of a chromosome sequence and *K* be the step size of the sliding window, then for a given interval size *S*, we produce the set of interval pairs *g_X_* × *g_Y_* = *G*, where

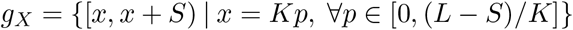

is the set of all *x*-axis image intervals (i.e. the intervals of size *S* at every *K^th^* position on the genome) and

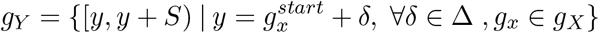

is the set of all *y*-axis image intervals, obtained by shifting the *x*-axis intervals by some *δ* (with 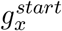 representing the location of the first base pair in *g_x_* and Δ a user-specified set of possible shift values). We use *δ* to find large events, where the end of the SV is further than *S* positions from the start of the *x*-axis interval. We can find smaller events by reducing the size of the intervals, *S*, to enlarge them in the image. In our experiments, we set *K* = 50kbp, *S* = 150kbp, and Δ = {0,75kbp, 150kbp}. Note that to detect events at any distance, including on different chromosomes, we can select interval pairs based on alignment information, e.g. by considering every pair of intervals that has a sufficient number of reads or read pairs that align to both. This optimization will be implemented in future work.

### SV confidence map regression

Given an *n*-channel image *I* generated for the genome interval pair (*g_x_,g_y_*), our neural network outputs a set of confidence maps *H* = {*h*_1_, *h*_2_,…, *h_T_*} corresponding to the *T* zygosity-aware SV types supported by the model. Each confidence map encodes the predicted location of the breakpoints of all the SVs of a particular type in the input. More specifically, let (*b_x_, b_y_*) be the coordinates of a given SV’s breakpoints in the genome (i.e. its start and end positions). If *b_x_* ∈ *g_x_* and *b_y_* ∈ *g_y_*, we can map these breakpoints to a keypoint 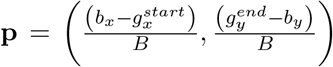 in *I*. If detected, this keypoint is sufficient to infer the genome coordinates of this SV within *B* base pairs by mapping its pixel coordinates back to genome space. Visually, the SV keypoint corresponds to the top-left corner of the square defined by the start and end coordinates of the SV on each axis of *I*.

We generate ground-truth confidence maps of size *w^H^* × *h^H^* to train the network as follows. For each SV, *v* (provided as a ground-truth BED or VCF file), overlapping *g_x_* and *g_y_* with a visible keypoint **p**^*v*^(i.e. 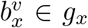 and 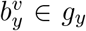), we add an unnormalized 2D Gaussian distribution peak centered around **p**^*v*^ to *h_t_*, where *t* represents the type of the SV. As a result, for each SV type *t* ∈ {1…*T*}, the values at 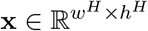 in *h_t_* are given by

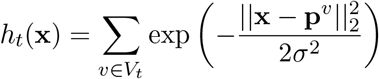

where *V_t_* is the set of all SVs in *I* of type *t*. The hyperparameter *σ* of the Gaussian kernel determines the spread of each keypoint peak and can be used to balance the ratio of foreground and background pixels. The confidence map size is determined by the stride hyperparameter *s*, which controls the ratio between the input image size and the confidence map size. In our experiments we generate confidence maps of size 64 × 64, given by *s* = 4 (with *σ* = 10).

### Network structure and training

Our deep learning model is a fourth-order stacked hourglass network based on the human pose estimation model proposed in [17, 18]. The network starts with a convolutional backbone module through which the image is fed prior to the four hourglass modules. The backbone consists of a 7×7 convolutional layer, a residual module, a max-pooling layer, and two additional residual modules. For input images of size 256×256, the backbone reduces the resolution down to 64×64 (our confidence map size). Each hourglass module consists of residual modules and max pooling layers to process the input features down to a low resolution, followed by nearest neighbor upsampling layers and skip connections to get back up to the output resolution. We perform intermediate supervision after each hourglass module, resulting in each hourglass module generating its own set of intermediate confidence map predictions from which we compute a loss. Stacking and intermediate supervision allow the network to repeatedly reassess its estimates and features at every scale. For more details on the HN architecture, please see [17, 18].

Our network was implemented in Pytorch. To train the network we used the Adam optimizer [39], a learning rate of 1*e*^-4^, and a batch size of 16.

### Focal loss

We use *focal L2 loss* adapted from [24] to compute the distance between the predicted and the ground truth confidence maps. Let 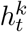 be the predicted confidence map of size *w^H^* × *h^H^* for SV type *t* by hourglass module *k* and let 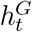 be the ground truth confidence map for this SV type, the focal L2 loss between these two confidence maps is defined as follows:

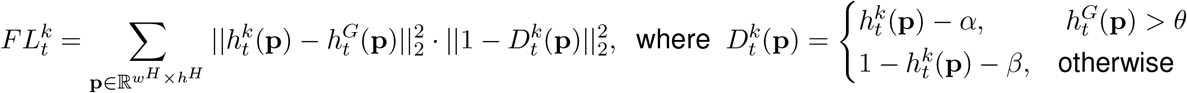

The hyperparameter *θ* is the threshold used to separate background and foreground pixels, while *α* and *β* are used to scale down the contribution of easy background and easy foreground pixels. The total loss for the stacked HN, summed over the four stacked hourglass modules and SV types, is then computed as:

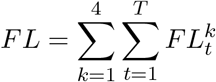

### Converting regressed confidence maps to final SV calls

Since the input images can contain multiple SVs of the same type, we detect all local maxima in each regressed confidence map using a maximum filter and thresholding (i.e. only values above a certain threshold are kept as candidates; we use 0.4 as the threshold in our experiments). Given the resulting set of peak keypoints, we perform 2D non-maximum suppression (NMS) by finding the SV breakpoint bounding boxes defined by each keypoint and filtering keypoints with conflicting or redundant bounding boxes. More specifically, let (*x*,*y*) be the coordinates of a candidate SV keypoint. The bounding box defined by this SV’s breakpoints is given by the following confidence map coordinates: [*x_min_* = *x*,*y_min_* = *y*,*x_max_* = *w* – *x*,*y_max_* = *h* – *x*]. For each pair of resulting bounding boxes (*M,N*), we compute the intersection over union, *IoU* = |*M* ∩ *N*|/|*M* ∪ *N*| and intersection over minimum, *IoM* = |*M* ∩ *N*|/min(|*M*|, |*N*|), metrics and remove the keypoint with the lower score if the *IoU* or *IoM* values are above a specified threshold (i.e. the boxes have substantial overlap).

In order to increase the accuracy of Cue’s breakpoint positions, we refine the location of the remaining key-points using higher resolution images “zoomed-in” around SV keypoints. During refinement, we extract a small patch of the initial image around each predicted keypoint and pass the patch through our model to obtain a higher resolution keypoint. To minimize the number of base pairs represented by each pixel, the input images are constructed using a smaller genome bin size (*B*), resulting in higher resolution.

Each refined SV keypoint coordinate (*x*,*y*) is then converted to genome breakpoint coordinates, with the *x* coordinate giving the start position of the SV and the *y* coordinate giving its end position (as previously described). Since each confidence map encodes only keypoints of SVs with a specific type (and genotype), we can directly determine the type and genotype of each SV call based on the index of the confidence map in which it was detected.

The above process produces a set of SV calls for each image, which are then collected and filtered using 1D NMS, wherein we compute the *IoU* and *IoM* metrics for the SV intervals on the genome to find and filter out near-duplicate or conflicting SVs. Since multiple images can capture the same part of the genome, and hence call the same SV, we need this step to remove such duplicate calls. Finally, our method can also be configured to filter out SVs falling into blacklisted regions of the genome (e.g. assembly gaps); however, this is not enabled by default.

The SV calling module was implemented to run in parallel on multiple CPUs or GPUs (by splitting the workload across chromosomes).

### Training data generation

To generate training data for our model, we simulated a human genome using SURVIVOR with 13,864 SVs of size 5kbp – 250kbp, consisting of homozygous and heterozygous deletions, tandem duplications, inversions, deletion-flanked inversions, inverted duplications, insertions, and translocations (note: insertions and translocations were included in the simulated genome but not labeled for training in the images). We have augmented SURVIVOR to also model specifically LINE-1 deletions, dispersed duplications, and inverted dispersed duplications (the latter two were not labeled as new event types in the images and instead served as negative examples). Additionally, we added small insertion and deletion events (of size 50bp - 1kbp) into this genome. We generated a paired-end Illumina short-read WGS dataset from this genome using DWGSIM and mapped the reads with BWA-MEM to obtain a BAM file with read alignments.

Given the generated BAM file and the ground-truth SV BED file produced by SURVIVOR, we generated an annotated training image dataset by: (1) scanning the genome using a sliding-window approach to produce genome interval pairs (we used intervals of size 150kbp for training and evaluation), (2) generating images from the alignments to each interval pair, and (3) annotating the resulting images with information that includes the SV type and breakpoint coordinates of each visible (or partially visible) SV, as well as genome intervals used to generate the image. This scheme resulted in 175,310 images with up to 6 SVs fully visible in the same image. In addition to this image dataset, we also generated 48,073 negative image examples using reads simulated directly from reference genome and 10,400 images with divergent read alignments (to model divergence, we simply downsampled reference-read alignments at random loci of the genome). Supplementary Figure A8 shows a high-level diagram of the in-silico training data generation process and a sample of produced annotated images.

### Evaluation metrics

In benchmarks where truthsets were available (i.e. synthetic and HG002 genomes), the SV calls and genotypes were evaluated using the benchmarking tool Truvari [31] to report the precision, recall, and F1 score as compared to the ground-truth SV callset. An SV call was considered a true positive (TP) if it had at least 50% reciprocal overlap and size similarity with an event in the ground-truth callset and was within 500bp of this event (Truvari parameters: `-pctovl 0.5 -pctsize 0.5 -refdist 500`); otherwise it was considered a false positive (FP). In order to evaluate performance in genotyping, we used the Truvari flag `-gtcomp`, which additionally compares the genotype of matching calls, such that TPs are considered only the calls defined above with matching genotypes, and FPs include the calls that either failed to match the genotype or the other constraints. Only SVs that passed all filters were considered (configured using the `-passonly` flag). When comparing SVs in a particular size range (configured using the `-sizemin` and `-sizemax` parameters in Truvari), we run Truvari twice to find the actual set of FPs as follows: (1) we match the evaluated callset against the ground-truth callset using the desired `-sizemin` and `-sizemax` thresholds and (2) we match the FPs obtained in (1) against all the calls in the ground-truth callset. This procedure allows us to find ground-truth matches that slightly fall outside the SV size thresholds (and are filtered by Truvari) – e.g. a 1000bp call that matches a ground-truth 999bp call would be considered a FP by Truvari with a `-sizemin` 1000 threshold.

For the CHM benchmark, the overlap between callsets was found using SURVIVOR [25], which merges SVs based on criteria such as the difference in SV breakpoint positions, event length, and event type. In our case, SURVIVOR was parameterized to merge calls within a breakpoint difference of 5kbp and with reported lengths of at least 1kbp. When identifying event overlaps across SV types (e.g. overlapping DEL and DUP calls), we considered two events as overlapping if the size of the overlap region was at least 80% the length of both of SVs.

## Data availability

The 60x HG002 Illumina WGS short reads, the 28x HG002 PacBio CCS reads, and the HG002 v0.06 truthset are available through the GIAB FTP data repository. In particular, short reads can be downloaded from ftp://ftp-trace.ncbi.nlm.nih.gov/giab/ftp/data/AshkenazimTrio/HG002_NA24385_son/NIST_HiSeq_HG002_Homogeneity10953946/NHGRI_Illumina300X_AJtrio_novoalign_bams/HG002.hs37d5.60X.1.bam, the PacBio CCS reads can be downloaded from https://ftp-trace.ncbi.nlm.nih.gov/giab/ftp/data/AshkenazimTrio/HG002_NA24385_son/PacBio_CCS_15kb/alignment/HG002.Sequel.15kb.pbmm2.hs37d5.whatshap.haplotag.RTG.10x.trio.bam, and the v0.06 truthset can be downloaded from ftp://ftp-trace.ncbi.nlm.nih.gov/giab/ftp/data/AshkenazimTrio/analysis/NIST_SVs_Integration_v0.6/HG002_SVs_Tier1_v0.6.vcf.gz. The CHM1 and CHM13 40x coverage Illumina WGS short reads can be downloaded from the ENA short read archive (ENA accessions ERR1341794 and ERR1341795, respectively). The CHM1 and CHM13 PacBio long reads can be obtained from the NCBI sequence read archive under accession numbers SRP044331 (CHM1) and SRR11292120-SRR11292123 (CHM13). The Huddleston et al. [21] CHM1 and CHM13 truthsets can be downloaded from http://eichlerlab.gs.washington.edu/publications/Huddleston2016/structural_variants. To obtain a single truthset, we merged the CHM1 and CHM13 VCFs using SURVIVOR and genotyped the calls accordingly (i.e. such that records reported in both CHM1 and CHM13 were labeled as homozygous, and records only reported in one of the two were labeled as heterozygous). To label duplications, we cross-referenced insertion calls with Supplementary Table 11 of [21], which separately reports which published insertion calls are duplications. All other data is available upon request.

## Acknowledgments

The authors would like to thank Harrison Brand, Chris Whelan, Xuefang Zhao, Aziz Al’Khafaji, and Mike Talkowski for useful feedback and discussions about the method. This study is supported by the Broad Institute Schmidt Fellowship Program (to V.P.). I.H is supported by start-up funds (Weill Cornell Medicine) and a NIGMS Maximizing Investigators’ Research Award (MIRA) R35 GM138152-01. We thank the WCM SCU and Cornell’s BioHPC Cloud for compute resources.

## Author Contributions

V.P. conceived the study and implemented the framework. V.P and I.H. designed the benchmarks. V.P. generated the training data and performed model training and evaluation across benchmarks. C.R. implemented benchmarking scripts to annotate and visualize SV callset consensus and performed the analysis for the CHM benchmark. F.C. evaluated SV candidate calls using long reads and deciphered the rearrangement patterns of several key mapping signatures. K.G. assisted with the interpretation of candidate SV calls and benchmark result analysis. D.M. produced the callsets of existing tools on benchmark datasets. I.H. provided GPU resources. V.P. wrote the manuscript. All of the authors revised the manuscript. V.P. supervised the study.

## Supplementary Tables

**Table A1:** Recall of complex INVDEL and INVDUP events. Matches were computed using Truvari by comparing only breakpoints (BP match) and both breakpoints and genotype (GT match). The total number of matches is reported along with the percentage shown in parentheses.

|  | INVDUP [704] |  | INVDEL [3078] |  |
| --- | --- | --- | --- | --- |
|  | BP match | GT match | BP match | GT match |
| Cue | 672 (95%) | 661 (94%) | 2707 (88%) | 2667 (87%) |
| Manta | 616 (88%) | 414 (59%) | 988 (32%) | 988 (32%) |
| LUMPY | 4 (0.5%) | 1 (0.1%) | 977 (32%) | 974 (32%) |
| DELLY | 647 (92%) | 429 (61%) | 998 (32%) | 991 (32%) |
| SvABA | 517 (73%) | 352 (50%) | 960 (31%) | 380 (12%) |

## Supplementary Figures

**Figure A1:**
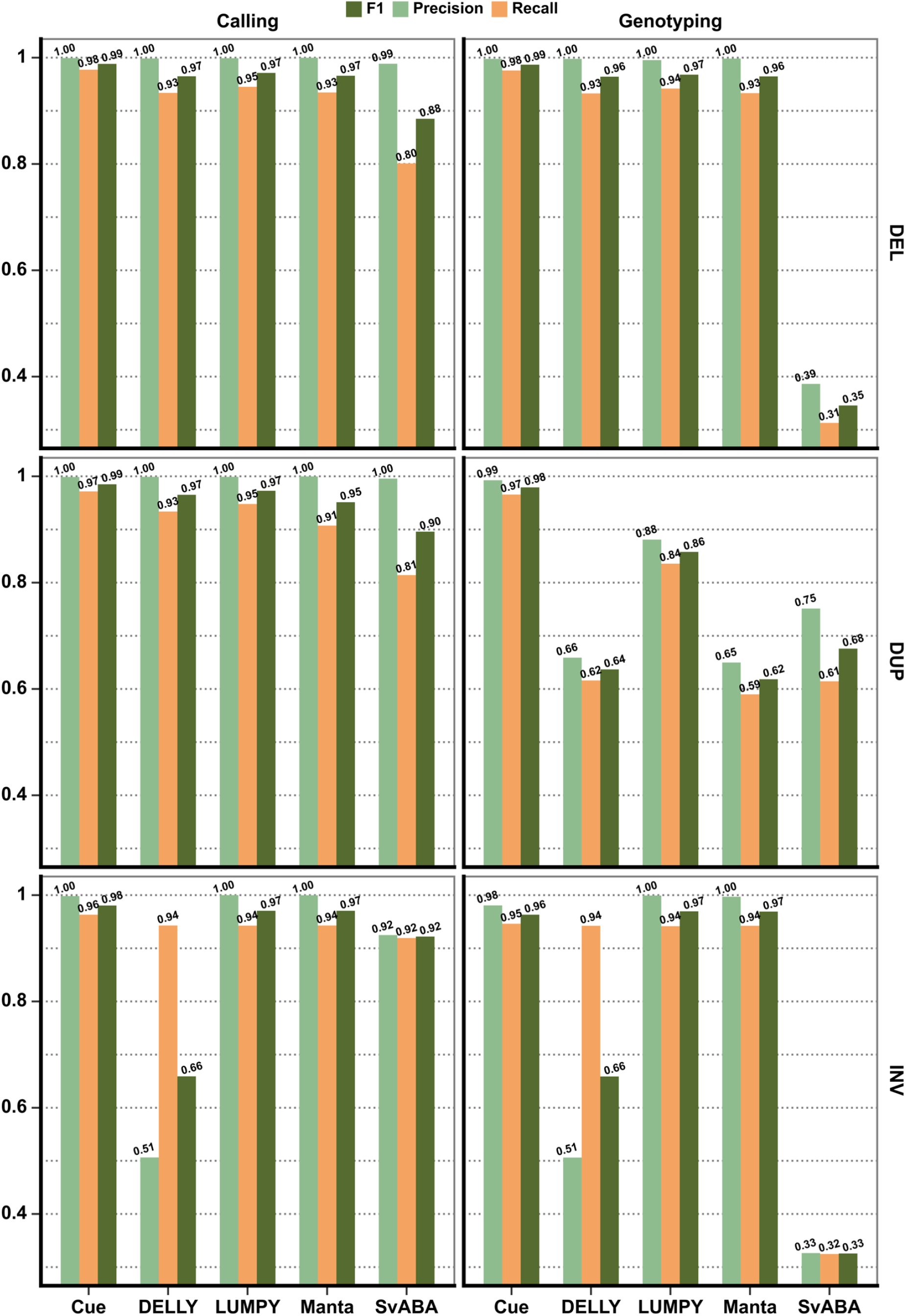
Performance evaluation of DEL, DUP, and INV calling and genotyping broken down by SV type on synthetic data.

**Figure A2:**
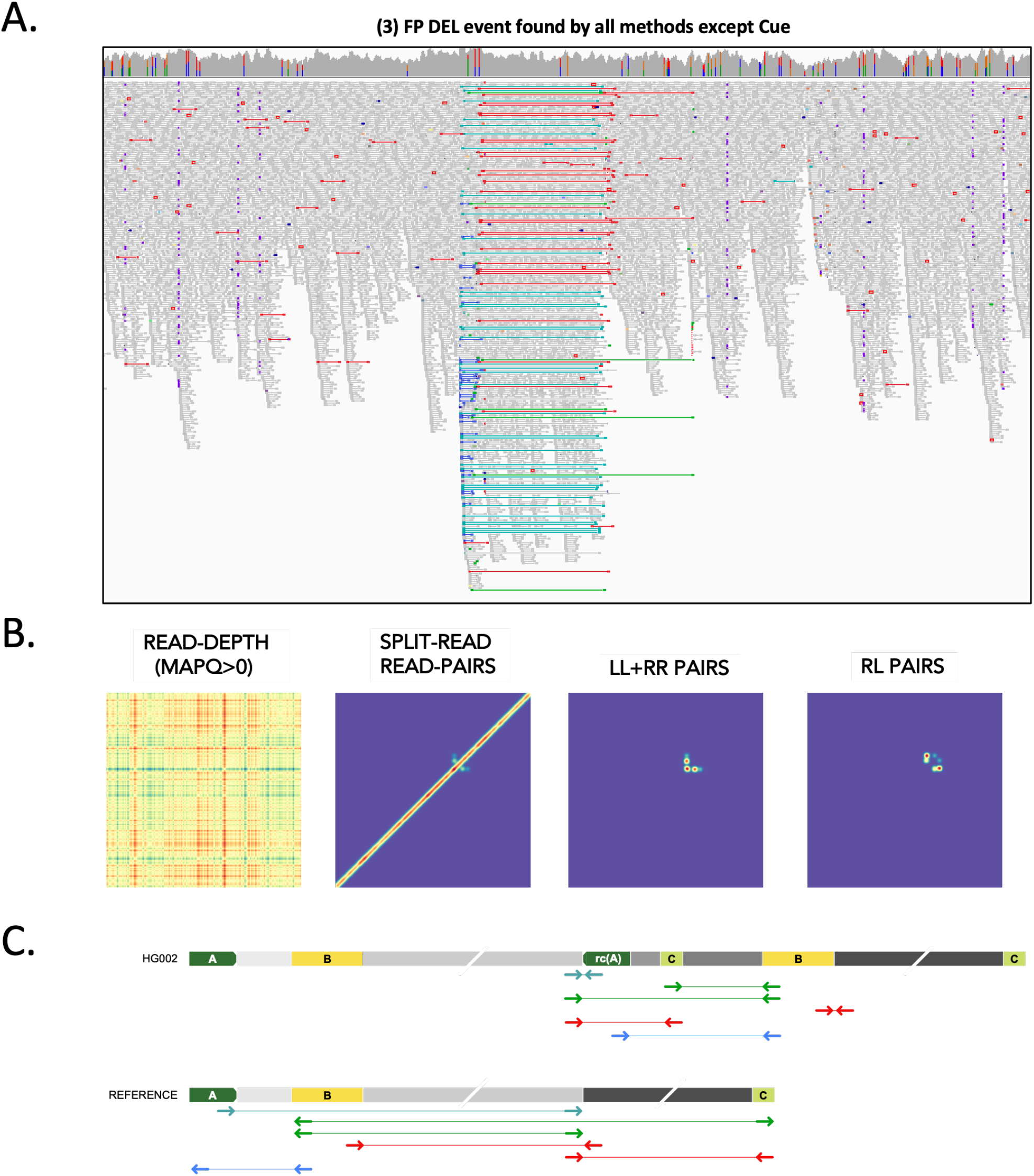
Analysis of a false positive HG002 deletion call generated by all short-read callers except Cue, using short and long reads. **A.** IGV plot showing short-read alignments around the call locus. Discordant read pairs mapped to the same strand (LL and RR mappings) are shown in light and dark blue (indicative of inversion), RL mappings are shown in green, and read pairs with a discordantly large insert size are shown in red. **B.** Cue-generated image channels depicting short-read signals; can be easily seen to be inconsistent with a valid DEL signature (see e.g. Figure 3C). **C.** One of the two haplotypes of HG002, reconstructed by de novo assembly of PacBio CCS reads, explains the main patterns of discordant pairs in panel **A** with two dispersed DUPs, one inverted dispersed DUP, and no DEL (the other haplotype is identical to the reference). Colored blocks labeled with letters are distinct short repeats. Gray blocks broken by diagonal lines are long sequences. *rc*(*A*) is the reverse-complement of *A*. Haplotypes were reconstructed and compared to the reference as follows. Let *W* be the sequence of the reference that covers the main patterns of discordant pairs in panel **A**. We built a joint de Bruijn graph (*k* = 87) on *W* and on the 190 CCS reads that have some alignment to *W*, we removed *k*-mers with frequency one, and we translated *W* and every read into a walk (which may contain cycles) in the graph.

**Figure A2:**
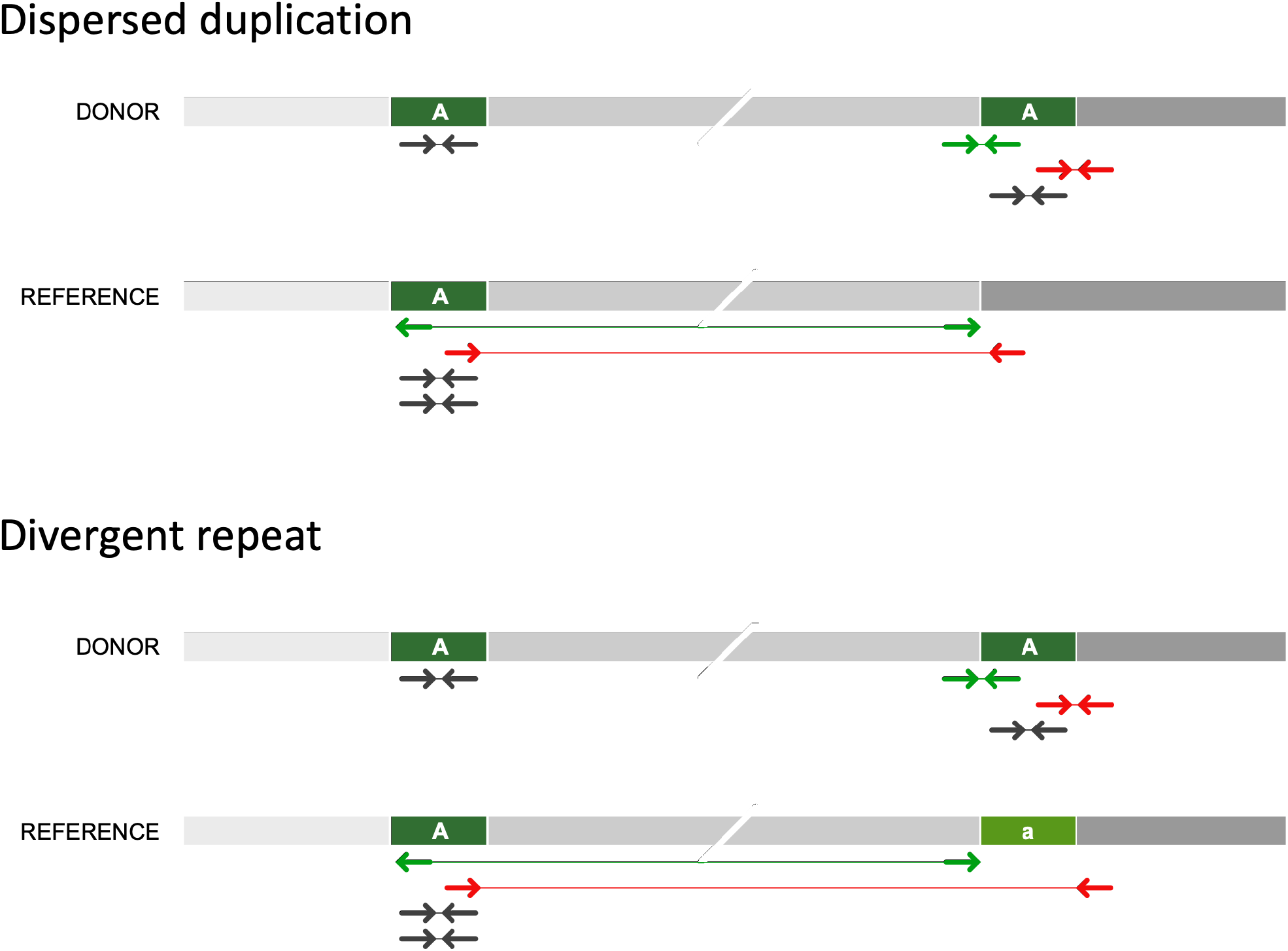
Schematic of a small dispersed DUP and a divergent reference repeat short-read mapping signature. Locus “A” is duplicated in the donor genome. Some read-pair fragments map discordantly in the RL orientation (green) or with a large insert size (red). Fragments internal to each copy of the donor map to the single copy of “A” in the reference genome, doubling its coverage. If the reference has a divergent copy of “A” (denoted as “a”), a gap in coverage will be observed at “a”.

**Figure A4:**
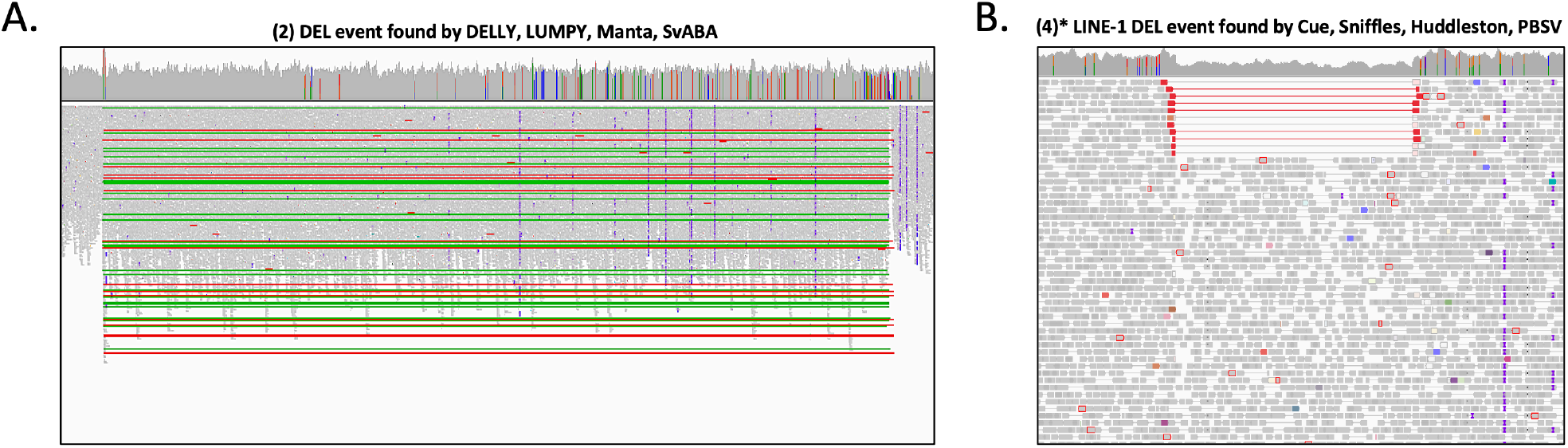
IGV plots of likely FP and TP calls in the CHM1+CHM13 benchmark. **A.** A DEL call reported by all short-read callers except Cue. **B.** A LINE-1 DEL event detected by Cue and all long-read callers.

**Figure A5:**
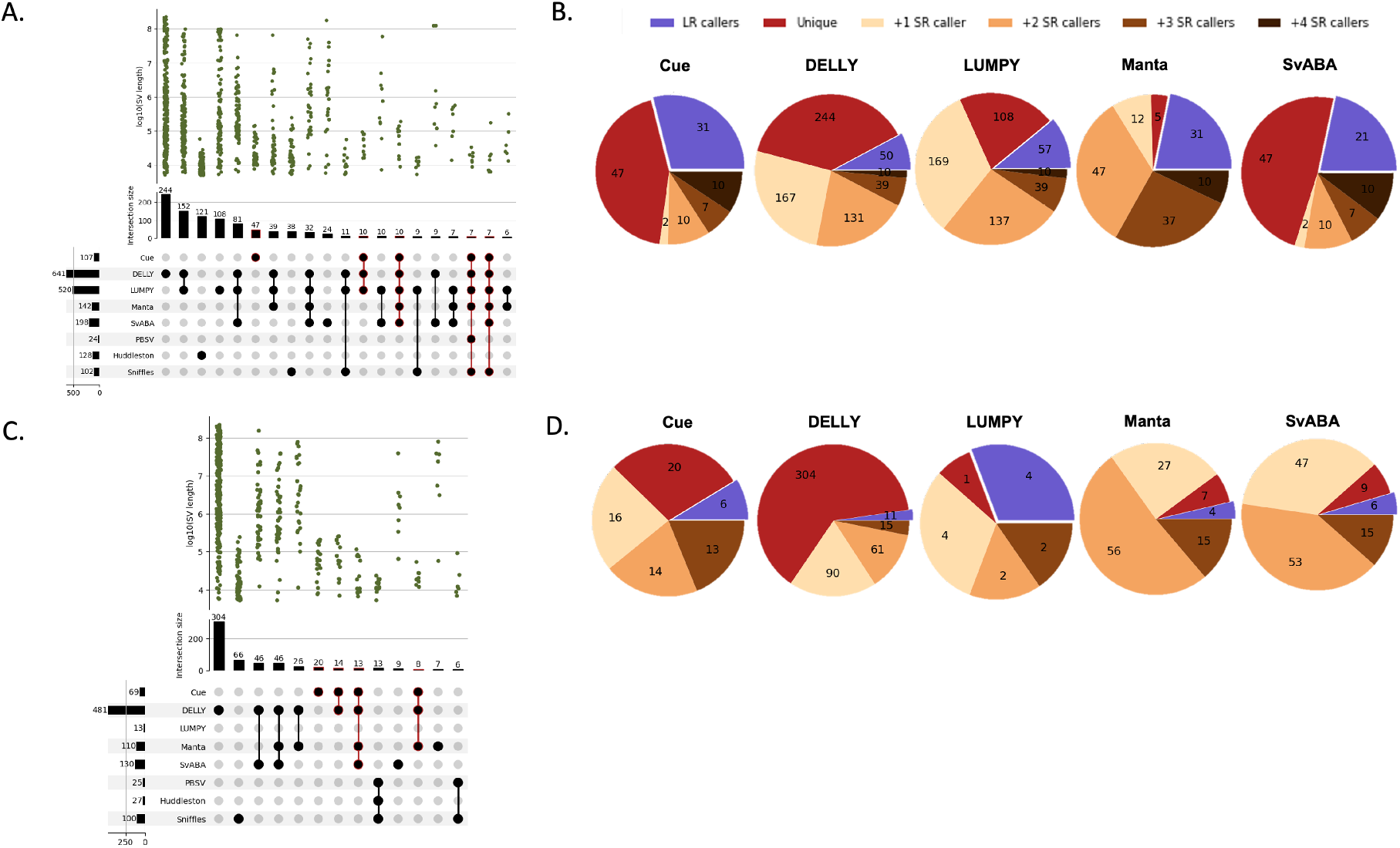
Performance evaluation of DUP and INV calling on the CHM1+CHM13 cell-line mix. **A.** Upset plot depicting DUP callset overlaps of short-read and long-read callers (only sets larger than 5 events are displayed for conciseness). **B.** Result breakdown of DUP calls by consensus with long-read and other short-read callers.**C.** Upset plot depicting INV callset overlaps of short-read and long-read callers. **D.** Result breakdown of INV calls by consensus with long-read and other short-read callers.

**Figure A6:**
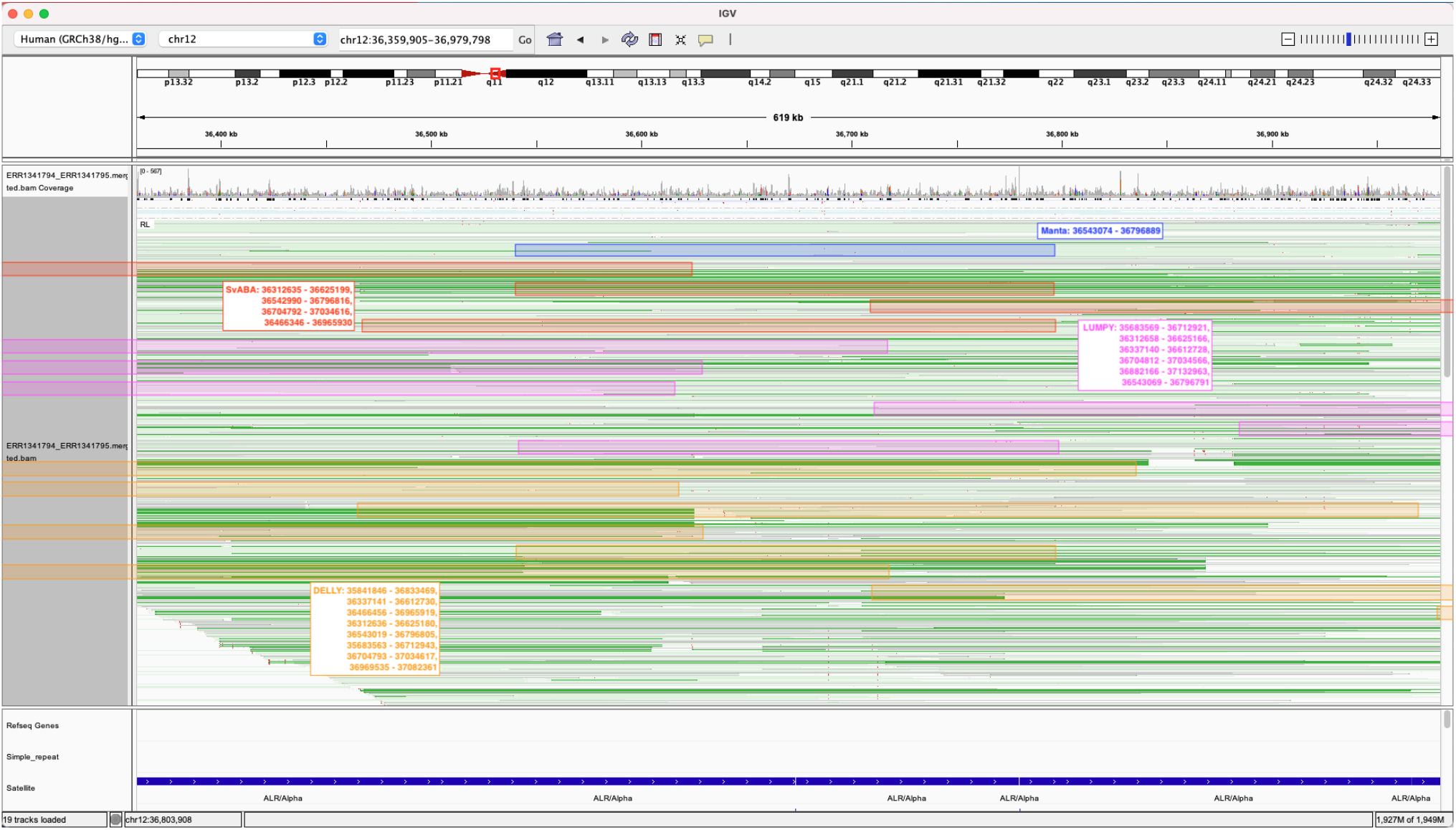
Manta, SvABA, LUMPY, and DELLY DUP calls in the chr12 centromere region of the CHM1+CHM13 cell-line genome mix.

**Figure A7:**
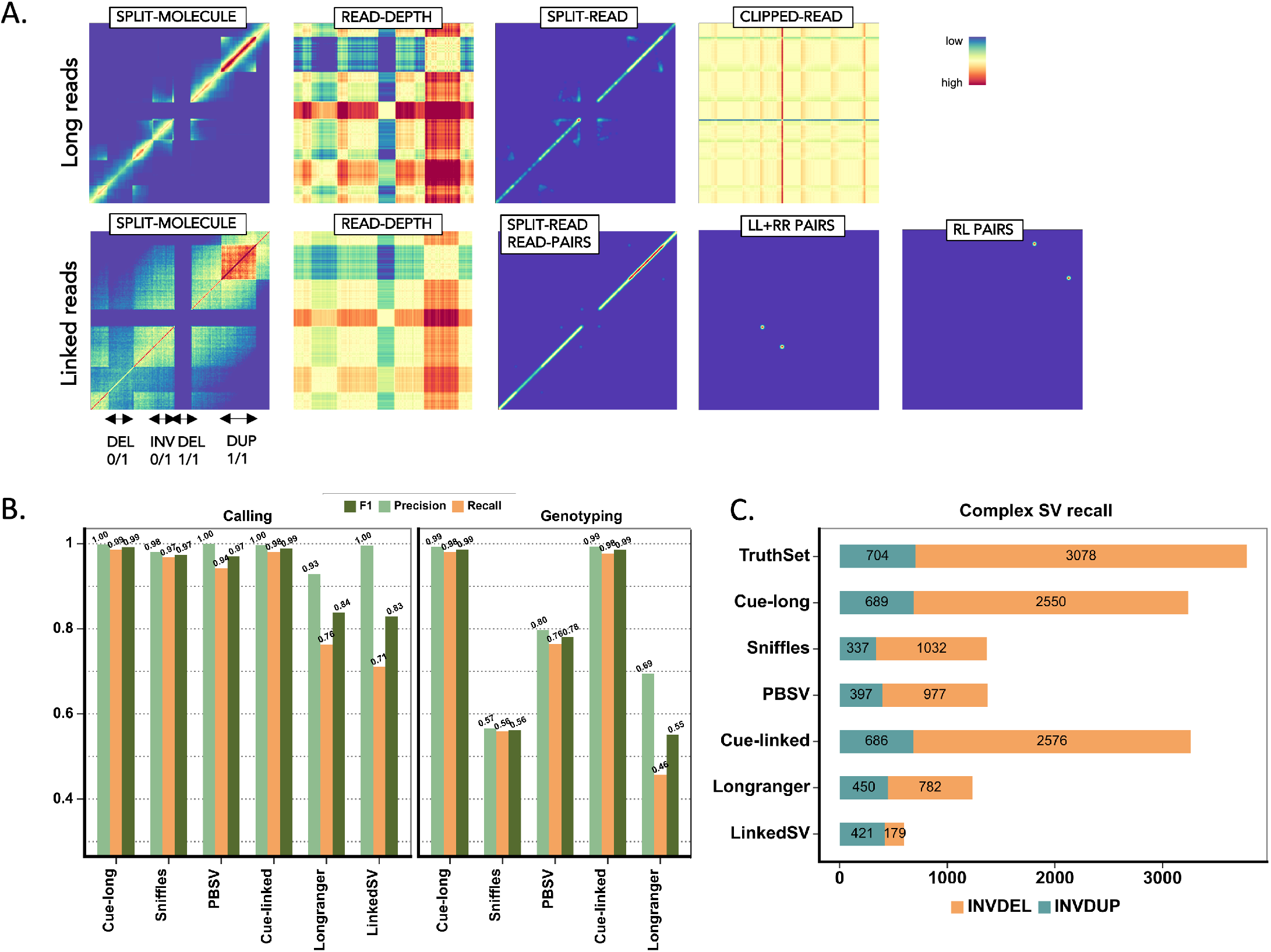
Extending Cue to long and linked read sequencing platforms. **A.** Image channels generated from synthetic PacBio CLR long reads and 10x Genomics linked reads, computed for an interval of the genome containing four different SVs (labeled along the *x*-axis; the same interval is assigned to both axes). **B.** Recall, precision, and F1 in DEL, DUP, and INV calling and genotyping. LinkedSV does not output SV genotypes and is omitted. **C.** Recall of two complex SV types (INVDUP and INVDEL) using long and linked reads.

**Figure A8:**
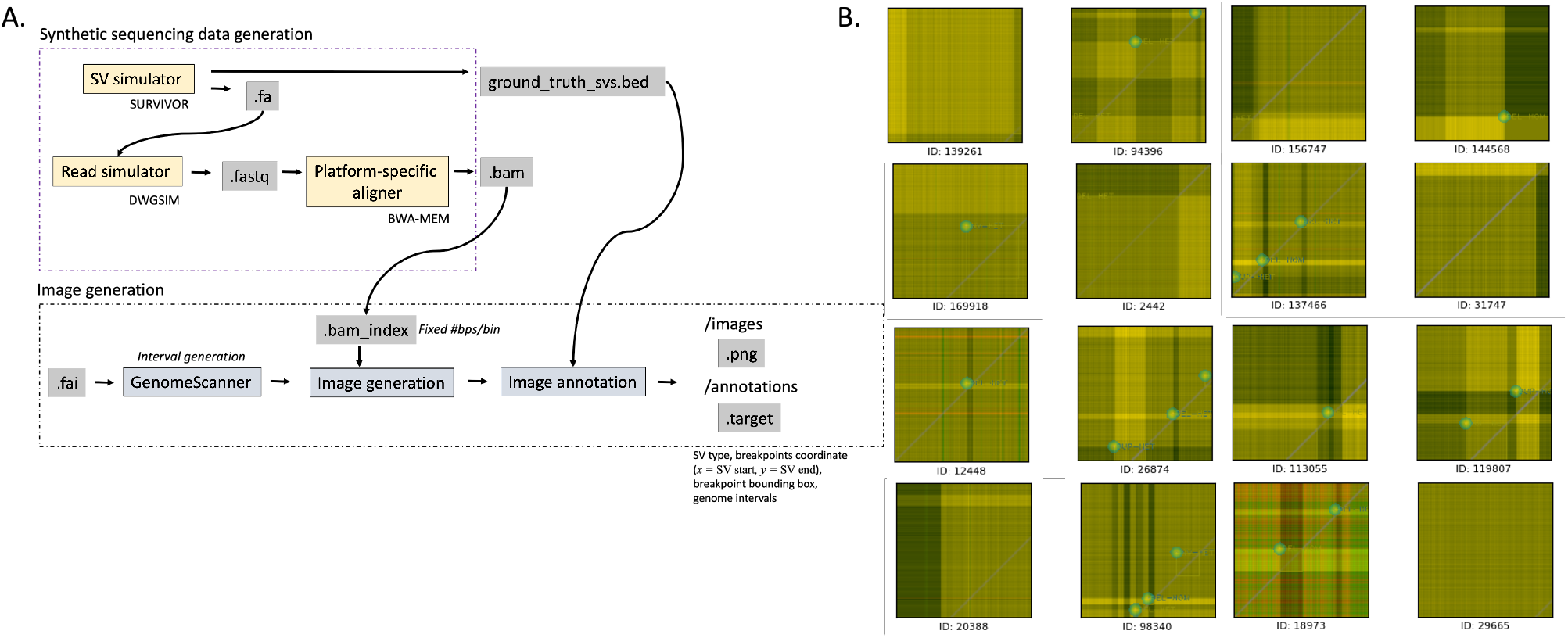
Training data generation. **A.** High-level overview of the in-silico sequencing and image data generation process. **B.** Annotated training examples (images are displayed for the figure using standard image visualization software based on only three of the constructed channels, including the read-depth channel).

## Supplementary Notes

### Software versions

The following versions of software tools were used in our benchmarks: BWA-MEM 0.7.15, SURVIVOR 1.0.3, Manta 1.6.0, LUMPY 0.3.1, svtyper 0.7.1, extractSplitReads_BwaMem 0.1.0, DELLY 0.9.1, SvABA, Sniffles 1.0.12, NGMLR 0.2.7, PBSV 2.6.0, pbmm2 1.3.0, IGV, samtools 1.9, bcftools 1.14, RCK, Longranger 2.2.1, LinkedSV commit 15259248d6f, and Truvari 2.1.

### Execution parameters

#### Manta

We followed the recommended steps to execute Manta by running the scripts `configManta.py, runWorkflow.py`, and `convertInversion.py` with default parameters.

##### LUMPY

We executed the following recommended workflow: (1) we extracted discordant reads using `samtools view -b -F 1294`, (2) we extracted split reads using the extractSplitReads_BwaMem tool, (3) we ran `lumpyexpress` with default parameters given the resulting files, and (4) we ran `svtyper` to produce a genotyped final VCF. We found that the LUMPY output often included numerous homozygous reference calls. We filtered such calls to improve LUMPY’s performance in the benchmarks (filtering consistently resulted in higher F1 scores).

##### DELLY

We ran the `delly` executable with recommended default parameters to call and genotype SVs and used `bcftools` to convert their output BCF files into VCFs.

##### SvABA

We ran the `svaba` executable with recommended default parameters and the `-germline` flag to call and genotype SVs (the flag was skipped in the subclonal benchmark). Since SvABA does not directly label the discovered events by their type, we used the tool RCK (namely, the `rck-adj-x2rck` and the `rck-adj-rck2x` scripts), which provides support to convert SvABA outputs to a standardized VCF that includes SV types.

#### Sniffles

On the CHM benchmark, we ran Sniffles with default parameters using the NGMLR alignments of the CHM PacBio reads as input (NGMLR was similarly executed with default parameters). For the synthetic benchmarks, we obtained results using both minimap2 and NGMLR inputs and reported the best of the two (namely, NGMLR for the complex SV benchmark and minimap2 for the basic SV benchmark). When computing the complex SV recall, we also allowed even the filtered SV calls made by Sniffles to match the ground truth events in order to increase its performance (since in this benchmark most of the reported Sniffles events were not annotated with PASS).

##### PBSV

We ran PBSV with default parameters using the recommended PBMM2 alignments as input (PBMM2 was similarly executed with default parameters and the `-median-filter` flag).

#### Longranger

We ran the `longranger wgs` pipeline with default parameters and `-vcmode=freebayes`. We combined the output `dels.vcf.gz` and `large_svs.vcf.gz` files. In order to correctly compare genotypes with ground truth using Truvari, all 1/0 entries were replaced with 0/1 entries.

#### LinkedSV

We ran the `linkedsv.py` script with default parameters and the `-germline_mode` flag. We found that LinkedSV outputs numerous near-duplicate calls, which can significantly lower its precision when benchmarking with Truvari. In order not to penalize these calls, we added the `-multimatch` flag in Truvari when evaluating LinkedSV, which allows for multiple reported calls to match the same ground truth call.

